# PyTME (Python Template Matching Engine): A fast, flexible, and multi-purpose template matching library for cryogenic electron microscopy data

**DOI:** 10.1101/2023.10.23.563472

**Authors:** Valentin J. Maurer, Marc Siggel, Jan Kosinski

**Affiliations:** European Molecular Biology Laboratory Hamburg, Notkestraße 85, 22607 Hamburg, Germany; Centre for Structural Systems Biology (CSSB), Notkestraße 85, 22607 Hamburg, Germany; Structural and Computational Biology Unit, European Molecular Biology Laboratory, Meyerhofstrasse 1, Heidelberg 69117, Germany

## Abstract

Cryogenic electron microscopy (cryo-EM) is a key method in structural and cell biology. Analysis of cryo-EM images requires interpretation of noisy, low-resolution densities which relies on identifying the most probable orientation of macromolecules in a target using template matching. Many method-specific template matching software exist for single-particle cryo-EM, cryo-electron tomography (cryo-ET), or fitting atomic structures into averaged 3D maps of macromolecules. Here, we report the Python Template Matching Engine (pyTME), a software engine that consolidates method-specific template matching problems. The underlying library provides highly efficient template-matching implementation and abstract data structures for storing and manipulating input and output data. It scales favorable to large datasets, both with multiple CPUs and GPUs, compared to existing software enabling template matching of even unbinned cryo-ET data in hours, which was previously nearly impossible due to technical restraints. Any hardware-specific optimization needed for dealing with large data is automatically performed to increase ease of use and minimize user intervention. The efficiency and simplicity of pyTME will enable high throughput mining of a variety of cryo-EM and ET datasets in the future.

## I. MOTIVATION AND SIGNIFICANCE

Cryogenic electron microscopy (cryo-EM) enables the study of biological specimens in a near-native state at up to single-digit Ångstrom resolution^1–3^. With advancements in sample preparation, detectors, and computational methodologies, the utility of cryo-EM has continuously increased, and different approaches for generating 3D density maps from 2D images have been developed^4^. Cryo-EM single-particle analysis (SPA) captures multiple 2D images of isolated and identical particles frozen in random orientations which are computationally aligned and averaged to generate a 3D reconstruction^3–5^. This method has been successfully used to solve the structures of individual proteins, such as small transporters in various conformations, up to large assemblies^6–8^. Cryoelectron tomography (cryo-ET) involves the collection of a series of 2D images (a tilt series) of the same region of a sample at different angles, which is aligned and back-projected to create a 3D reconstructed volume^9,10^. Cryo-ET enables the exploration of the spatial organization of macromolecules *in situ* in their native environment^11–14^. Individual subtomograms can be further averaged to yield high-resolution averages^9,15^.

Template matching is a crucial method for analyzing EM data and finds applications in localization, reconstruction, alignment, averaging, and classification. Conceptually, it quantifies the similarity between a template and a target and can be used to identify an orientation of the template that has maximal similarity to the target. A common similarity metric is cross-correlation, but recent years have also seen deep learning-based approaches^16,17^. In the context of localization, template matching is referred to as particle picking on 2D images from SPA, fitting atomic structures to low-resolution density maps, and template matching on 2D tilts or 3D reconstructions in cryo-ET^18–20^. Similarly averaging, or sub-tomogram averaging in the case of 3D tomograms, relies heavily on template matching to ensure correct alignment of the input data.

Several challenges are associated with template matching. The data is often noisy, prone to contrast variations, and the resolution is limited. Additionally, macromolecules can display a high degree of conformational variability, introducing additional uncertainty and decreasing resolution. Template matching is also a computationally challenging task, as the complexity for a *N* -dimensional hypercube with edge length k is at best 𝒪(*k*^*N*(*N−*1)^ log(*k*^*N*(*N−*1)^)) (see II A). Recent work showed the potential of template matching for comprehensive annotation of cells but also noted these severe computational restraints by existing software^21^.

Here, we present pyTME, a unified template matching engine that provides abstract data structures, extensive preprocessing methodologies, and efficient data type- and dimension-agnostic template matching methods. pyTME can be combined with the classification, averaging, and reconstruction procedures of existing software as a modular template-matching solution. We show that pyTME is easy to use, has higher test coverage than comparable tools, and excels in benchmarks.

## II. SOFTWARE DESCRIPTION

pyTME is a Python library that generalizes template matching, implements efficient data structures and provides preprocessing methods for cryo-EM data. pyTME gives practitioners access to high-performing versatile template matching and enables computationally advanced users to build their own software using pyTME’s API. pyTME is rigorously tested with code coverage of 95% (3,933 of 4,152 statements tested), which is drastically higher than for comparable software, such as PyTom^22^ with a test coverage of 18% (5,822 of 31,479), PowerFit^23,24^ with 24% (425 of 1,754) and STOPGAP^25^ as well as colores (SITUS)^26,27^ with 0% as no tests were available. pyTME’s high code coverage ensures correct implementation of existing code and seamless integration of novel features. pyTME operates on abstract class concepts, which are shown in an overview in Figure 1 and elaborated in more detail in the following.

**FIG. 1.**
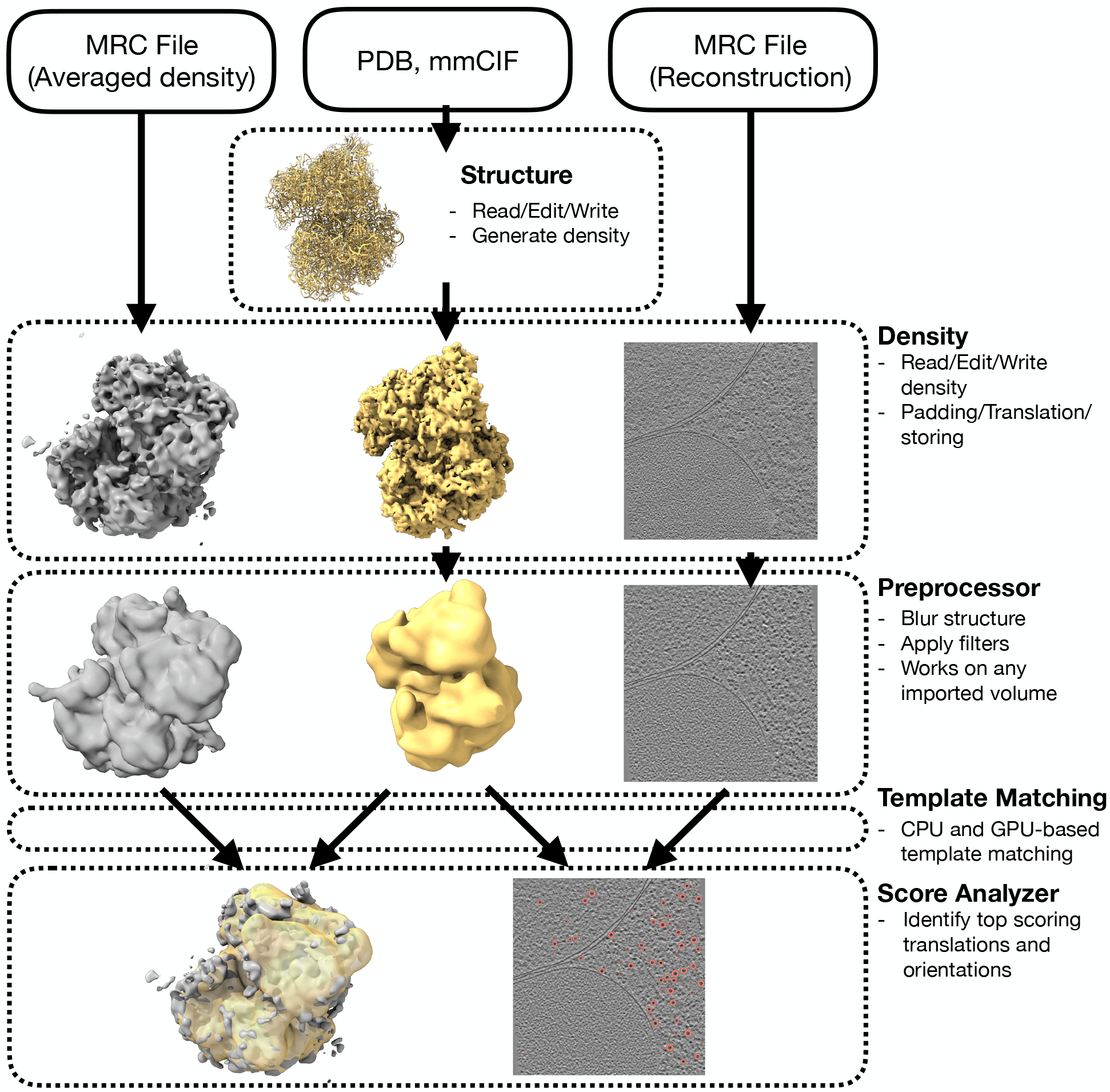
Overview of pyTME’;s main features and example workflows. Shown in the Density row is an EM map (left, EMD-3228, ref. 28) and atomic structure (center, PDB-ID: 6S47, ref. 29) of the *S. cerevisiae* 80S ribosome, as well as an *S. pombe* tomogram (right, EMPIAR-10988, TS 037, ref. 16).

### A. Software architecture

#### Structure

The `Structure` class represents macromolecular 3D atomic structures. The `Structure` class is instantiated from atomic structures files in Protein Data Bank (PDB)^30^ or mmCIF format. Its attributes are arrays that correspond to the atomic coordinate section of a PDB file and a header dictionary that stores structure meta-data. Once instantiated, the `Structure` class defines a range of operations. Subsetting enables selecting subsets of residues or chains from the structure, rigid transformations, i.e., translations and rotations, as well as comparison and alignment with other `Structure` instances. Including these standard functionalities avoids users needing additional tools for processing. The class instance of a structure can be either written to disk or converted to a density representation which is a prerequisite for the following template matching procedure. pyTME implements standard methods of converting atomic structures into densities based on atomic number, van der Waals radii, or atomic scattering factors^31,32^.

#### Density

The Density class serves as an abstract representation of densities and operates on *N* -dimensional arrays with defined sampling rate and origin, but for the reported cases we use only *N* = 3. The `Density` class can be instantiated from a variety of file formats such as CCP4/MRC, atomic structures in PDB/mmCIF format, or `Structure` class instances. The `Density` class defines a range of operations on its internal data array such as trimming, padding, and resampling. In addition, the `Density` class defines rigid transforms, coordinate system transforms, and alignments with other `Density` instances. `Density` class instances can be written to disk in various formats such as CCP4 or MRC for later use.

#### Preprocessor

The `Preprocessor` class defines filtering operations and applies them to arrays such as the ones contained within Density class instances. These filtering operations include classic filters known from image processing such as Gaussian, mean, and bandpass filter, as well as EM-specific filters, non-local filters, and user-supplied kernels. The `Preprocessor` class also enables the creation of wedge-masks for cryo-ET data^22,25^ and Fourier cropping to reduce the size of the input array^25^.

To simplify the usage of the `Preprocessor`, we provide a Napari^33^ plugin (Fig. S1), to access many of the functionalities via a graphical user interface. File formats supported by the Density class can be directly imported, inspected processed, and saved for subsequent use. After selecting a filter and parameter that leads to a transformation adequate for the user, the filter configuration is exported as YAML files for reproducibility and processing of larger datasets. In addition to filters, the Napari plugin also enables creating masks for template matching, provides shortcuts to automatically define suitable masks, and can be used to evaluate template matching results.

#### Template matching

Template matching in pyTME can be performed using two main strategies. Either all translations are exhaustively sampled together with a specific angular sampling density, or an underlying scoring function is numerically optimized. The following outlines the implementation details of both strategies.

#### Exhaustive template matching

Exhaustive template matching routines are designed in a modular fashion and require setup and a scoring function. The setup function pre-computes and caches operations that will be subsequently used and enables sharing them in memory between processes. The scoring function defines the exact procedure used to compute template-matching scores. This design has several key advantages: 1.) RAM usage is significantly reduced by avoiding data copies. 2.) New scores can be easily implemented by reusing existing code. 3.) Optimal parallelization schedules can be computed ahead of time (section II C). Exhaustive template matching scores shipped with pyTME are outlined in Tab. I and further described in the supporting information.

#### Non-exhaustive template matching

Non-exhaustive template matching uses numerical optimization to determine the translation and rotation of a template that maximizes an underlying scoring function. Depending on the topology of the sampled score space, non-exhaustive template matching can identify different maxima depending on the initialization conditions. Nonexhaustive template matching is primarily used for refinement or if the template occurs once or a few times, e.g. an atomic structure and a low resolution EM map.

pyTME uses the basin-hopping and differential-evolution methods from the SciPy library^38^ for global optimization with linear constraints. Since the optimal optimization scheme ultimately depends on the topology of the score space, other optimization schemes can be easily implemented by inheriting from existing abstract base classes.

**TABLE I.** Exhaustive template matching scores implemented in pyTME. Equations are listed in the Supporting Information.

| Score | Equation | Reference |
| --- | --- | --- |
| Cross-Correlation | 3 | 34 |
| Normalized Cross-Correlation | 4 | 34 |
| Laplace Cross-Correlation | 5 | 26,35 |
| Correlation About Mean | 6 | 32 |
| Masked Cross-Correlation | 8 | 36 |
| Fast Local Correlation | 7 | 22,25,37 |

Non-exhaustive template matching supports all scoring functions outlined in the previous section as well as additional scores that are outlined in Tab. II further described in the supporting information^39^.

**TABLE II.** Non-exhaustive template matching scores implemented in pyTME according to Vasishtan and Topf 39 . Equations are listed in the Supporting Information.

| Score | Equation |
| --- | --- |
| Partial Least Square Difference | 9 |
| Normal Vector Score | 10 |
| Envelope Score | 11 |
| Chamfer Score | 12 |
| Mutual Information Score | 13 |

#### Score analyzer

While non-exhaustive template matching results in a single maximum that requires no further analysis, the output of exhaustive template matching on 3D data is a 6D score space. Most tools reduce this to a 3D score space by aggregation^22–27^, so that only the maximum score for each translation over all rotations is saved. Unlike other tools, pyTME provides full flexibility over the analysis of scores through a dedicated `Score analyzer` class. pyTME is shipped with a variety of analyzers that can reproduce the aggregation behavior described above, generate peak lists, or perform clustering within the six-dimensional score space.

### B. Flexible backend

pyTME implements a flexible computation backend that allows for hardware-agnostic parallel code execution on CPU and GPU using a best-of-breed approach. In practice, flexible backends are implemented by wrapping array backends such as NumPy, PyTorch, and CuPy into a common NumPy-like syntax^40–42^. Each backend is required to define template matching related operations such as rigid- and Fourier transforms, as well as conversion methods that allow for moving array objects across devices.

The flexible backend allows for modifying particular components of the computational execution pipeline such as the used array type or Fourier transform strategy without having to modify the underlying template matching code. In combination with a rigorous abstract backend specification, the flexible backend ensures API stability while providing full flexibility. By following the abstract backend specification, new backends can be easily integrated into pyTME without knowledge of the underlying template matching operations.

### C. Automated RAM optimization

Using more memory allows for shorter runtime by avoiding overhead from inter-process communication and repeated initialization of temporary data containers. However, in practice, only a limited amount of memory is available per CPU/GPU on a compute cluster. Large input data such as tomograms are typically split into subsets for template matching that are then processed in parallel to achieve a balance between memory usage and runtime. This operation is supported by PyTom^22^ and STOPGAP^25^, but defining a suitable splitting strategy is a challenging task and typically has to be performed manually by the user.

To optimally use the available resources, pyTME internally computes how the input should be split and how many CPUs/GPUs should be allocated to processing each data chunk. To achieve this, we implemented methods that estimate the memory usage for all possible cases. Subsequently, pyTME determines the splitting strategy that minimizes overhead from the parallelization framework, subject to constraints on available memory and compute units.

This RAM optimization procedure hides the obfuscated and time-consuming task of determining a splitting strategy from the user, enables better utilization of available compute resources, and allows pyTME to easily scale to large datasets even in the hands of less experienced users of compute clusters (Fig. S2).

## III. ILLUSTRATIVE EXAMPLE

The following outlines how to use pyTME to perform template matching and recapitulate Fig. 1.

**Listing 1.**
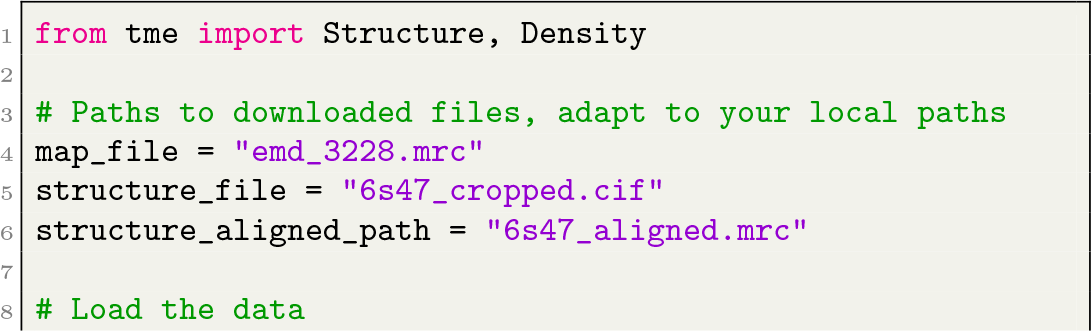
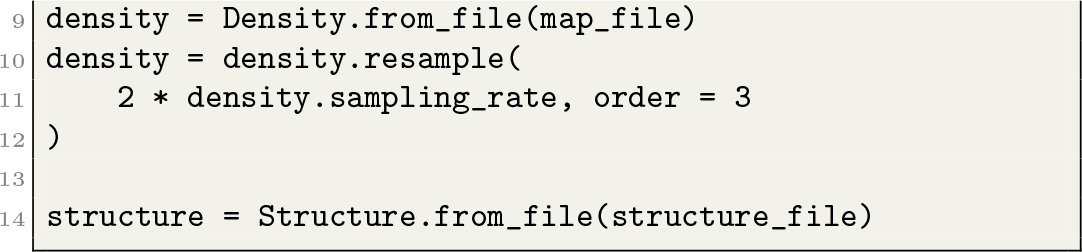
Load density and structure

The from file methods of the density and structure class are used to load the input data. They create class instances of the respective given type (Listing 1) and internally determine an appropriate parser for the data type. The data is resampled to twice the voxel size using cubic spline interpolation, in turn reducing the amount of data.

**Listing 2.**
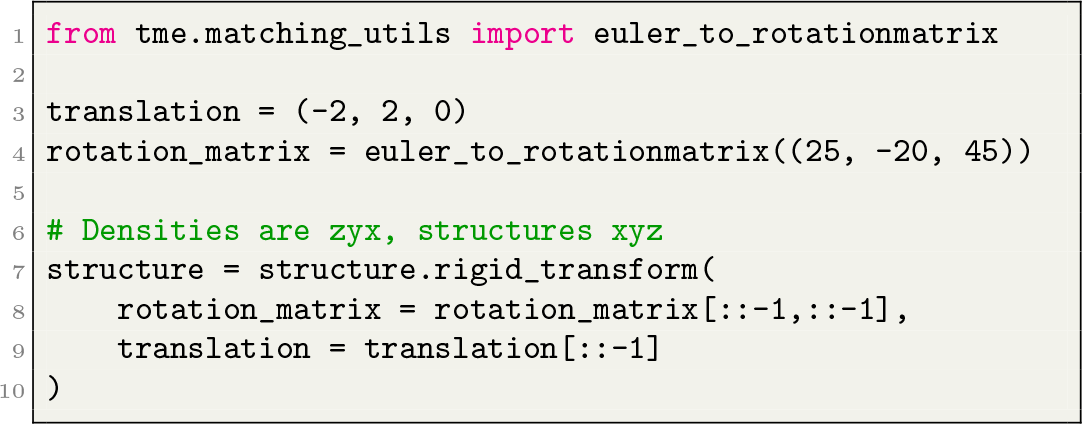
Rotate and translate structure

Since the example structure is already correctly oriented within the density, we perform a rigid transform to obtain an erroneous orientation (Fig. 1), aiming to recover the correct initial orientation using template matching later (Listing 2).

**Listing 3.**
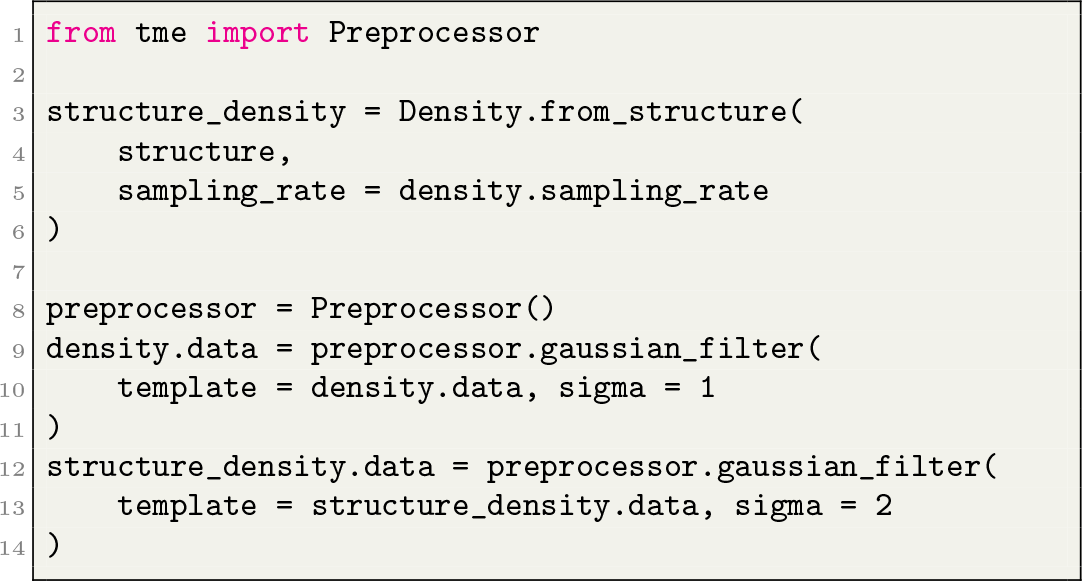
Preprocess the structure

Next, we convert the structure instance to a density and apply a Gaussian filter to the internal data arrays of both `Density` instances (Listing 3). At this point, the data should look like in the `Preprocessor` row in Fig. 1.

**Listing 4.**
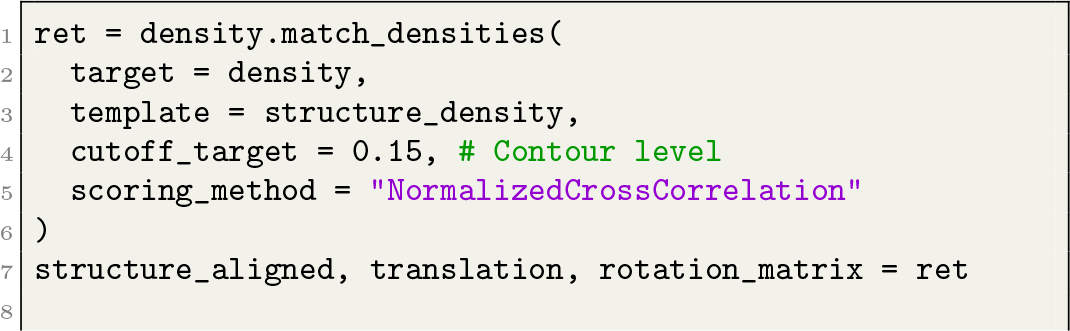
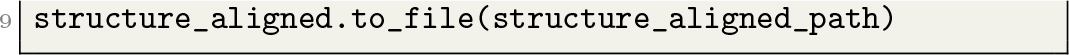
Non-exhaustive template matching

Listing 4 shows non-exhaustive template matching using normalized cross-correlation (equation 4). The output is a `Density` instance representing the aligned structure, the recapitulated translation, and the corresponding rotation matrix. The to file method saves the aligned `Density` instance to disk as CCP4/MRC file. This example recapitulates Fig. 1 (left). No further tools are required and the code can be easily adapted for other usage examples.

To showcase pyTME’s command line interface, we perform exhaustive template matching between a tomogram and an averaged EM density map: The output file match-ing result.pickle contains scores and rotations used to obtain high-scoring candidates analogous to the output of other template matching software^22–25^. In Fig. 1 we used an angular sampling rate of 7, which requires more computational resources. High-scoring translations and their corresponding rotations can be extracted as follows:

**Listing 5.**
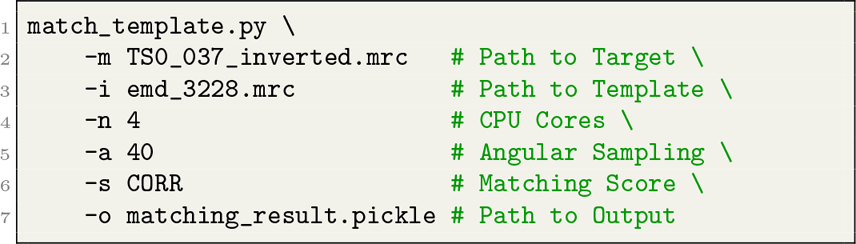
Cryo-ET template matching

**Listing 6.**
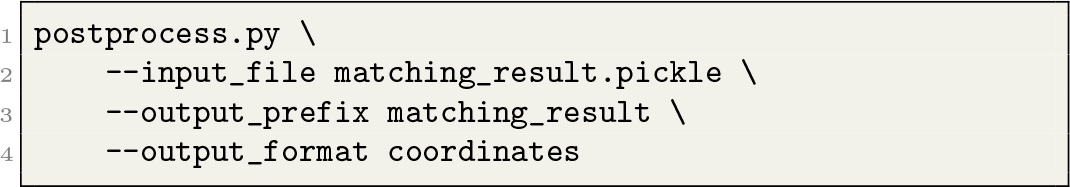
Score analysis

The resulting tab-separated file, contains eight columns, and each row corresponds to a translation and rotation of the template resulting in a given score and additional analyzer-specific information. The information contained within can be used by pyTME or other software in subsequent analysis. Postprocessing supports further functionality which is described extensively in the documentation.

## IV. PERFORMANCE AND SCALING BENCHMARKS

We benchmarked pyTME’s runtime performance against Situs (3.1)^26,27^, PowerFit (2.0.0)^23,24^, PyTom (1.0)^22^, and STOPGAP (0.7.1)^25^ (Fig. 2). Situs and PowerFit are tailored for fitting atomic structures into EM maps, while PyTom and STOPGAP focus on particle picking from tomograms. STOPGAP, PyTom, and Situs provide additional features that are beyond the scope of pyTME, which focuses on providing efficient data structures and template matching. Tools were containerized according to their respective manuals, if applicable. STOPGAP and the GPU version of PyTom were run uncontainerized. When running PyTom on the CPU, the target was split once along the second and third axis, because PyTom would otherwise exceed the RAM limit of 350 GB. CPU benchmarks (Fig.2 A and B) were performed on compute nodes equipped with AMD Epyc 7502 CPUs and 3,2 MHz DDR4 memory, while GPU benchmarks (Fig. 2 C) were performed with NVIDIA A100 GPUs. pyTME was run using mixed precision which is not yet supported by PyTom, but improves performance by 10-15% and potentially more on optimized hardware.

**FIG. 2.**
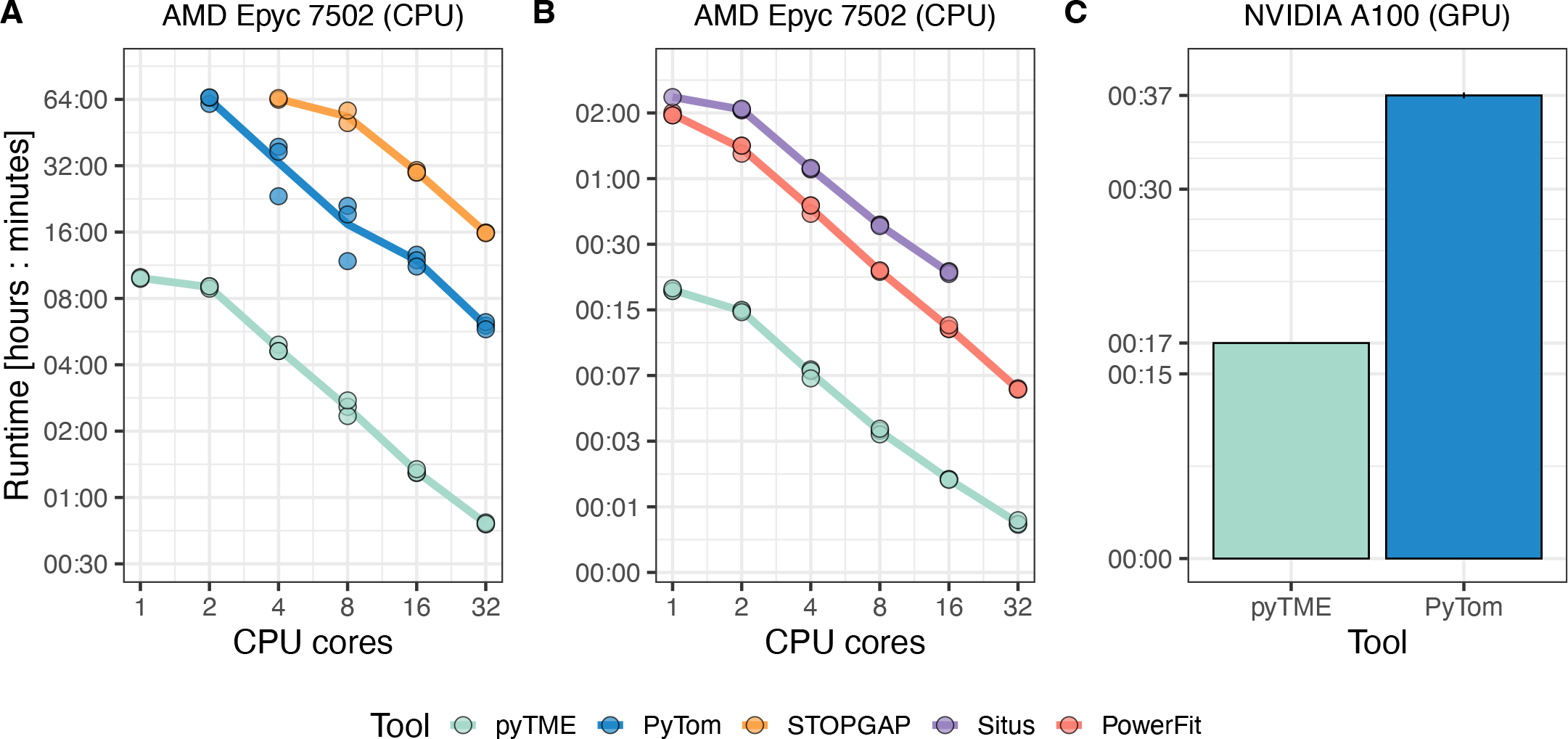
Runtime comparison of tools for particle picking (CPU: **A**, GPU: **C**) and fitting (CPU: **B**). For **A** and **C**, we used a tomogram as a target (EMPIAR-10988 43, TS 037, ref. 16 ) and a ribosome as template (EMD-3228; ref 28) as shown in figure 1. The dimensions of the target were 500, 960, and 928, and the template 51, 51, and 51. For **B**, we used a reconstructed cryo-EM map of the *E. coli* 30S ribosome subunit as the target (EMD-8621, ref. 44) and its atomic structure as the template (PDB-ID:5UZ4, ref. 44). The dimensions of the target were 220, 220, 220 and 124, 153, 120 for the template. Each tool sampled 1,992 rotations, apart from Situs, which sampled 1908. In **C** pyTME sampled 27,672, and PyTom 15,192 rotations, and the runtimes were normalized based on this ratio. Shown in **A** and **B** are individual trials and their average. **C** shows average runtime across three trial and standard deviation.

pyTME outpaces the tested software packages, achieving a speedup of over ten times in template matching for tomograms (Fig. 2 A) and over five times for electron density maps (Fig. 2 B). This can be similarly seen on GPU, albeit then the speedup over PyTom is lower (Fig. 2 C). pyTME achieves better performance by caching Fourier transforms, pre-allocating and sharing arrays in memory across processes as well as by using in-place operations. Software libraries implementing GPU operations such as CuPy^41^ used in pyTME and PyTom cache a majority of operations internally. This behavior of CuPy causes PyTom to also benefit from features that resulted in the outperformance of pyTME in Fig. 2A and B.

Runtimes are primarily determined by the number of Fourier transforms used to compute the scores. STOP-GAP and PowerFit compute different scores that require three instead of one Fourier transformation per rotation. Therefore, their runtime can be reduced by a factor of three for alignment with other tools. However, we did observe that PowerFit used twice the amount of RAM used by pyTME for comparable scores. Padding strategies are essential to numerical stability in Fourier transforms and the reduction of boundary effects but have less effect on runtime. Therefore, STOPGAP’s runtime can be reduced by an additional 10%. Unlike other tools, pyTME gives the user full flexibility over the padding approach.

## V. IMPACT AND CONCLUSION

pyTME brings template matching to users and developers in a unified data type-agnostic engine for easier accessibility and extendability together with extensive tutorials, documentation, and scripts. This removes the necessity to maintain multiple libraries for various applications which conceptually all require the same framework. Developers can expand on the data-type agnostic framework to easily test and develop new methodologies. Moreover, pyTME gives access to a broad range of scoring methods and sampling strategies, which were previously limited to either template matching in cryo-ET data or matching atomic structures. pyTME scales efficiently to large datasets such as tomograms at low binning (Fig. S2), and can be readily integrated into existing workflows.

pyTME outperforms existing software in terms of run-time, making it feasible to analyze larger datasets. The performance will allow the processing of larger and more complex data sets. With pyTME, tomograms can be annotated at low binning (Fig. S2) or datasets consisting of many files can be analyzed faster. Efficient template matching is also an essential prerequisite to generating training data for machine learning approaches, which require annotated datasets as input^16,17^. Higher resolution will eventually also allow the identification of smaller macromolecules or variations of templates indicative of different conformational or rare states. Therefore, the improvement in speed allows for faster turnover and better mapping of many different states in tomography. In the future, we envision that improvements in scoring and analysis will also further enhance the performance of template matching.

## VI. CODE AVAILABILITY

pyTME is available from https://github.com/KosinskiLab/pyTME. Code used to generate the benchmarks in this study is available from https://github.com/KosinskiLab/pytme_publication.

## VII. CONFLICT OF INTEREST

The authors declare no conflicts of interest.

## ACKNOWLEDGMENTS

We thank the EMBL IT and HPC resources for providing essential computational infrastructure. We thank Rasmus K. Jensen (EMBL, Heidelberg) for providing reconstructed tomograms and template at different binning. VM and JK acknowledge funding from the CSSB flagship project Plasmofraction. MS acknowledges support from a research fellowship from the EMBL Interdisciplinary Postdoc (EIPOD) Programme under Marie Curie Cofund Actions MSCA-COFUND-FP (grant agreement number: 847543).

## Notes

https://github.com/KosinskiLab/pyTME

### Competing Interest Statement

The authors have declared no competing interest.

