## Supporting Information for "PyTME (Python Template Matching Engine): A fast, flexible, and multi-purpose template matching library for cryogenic electron microscopy data"

Exhaustive template matching, as implemented in pyTME, uses cross-correlation to determine the similarity between a target and a template:

$$(f * g)[n] = \sum_{m=-\infty}^{\infty} f[m]g[n+m] \tag{1}$$

where $*$ denotes the convolution operator, f and g - the target and the template sampled at spatial location $[x]$. The computational complexity of this operation on two cubes with edge length N is $\mathcal{O}(N^6)$, but it can be reduced to $\mathcal{O}(N^3\log(N^3))$ through the application of the cross-correlation theorem:

$$(f * g)(t) \longleftrightarrow \mathcal{F}^{-1}(\mathcal{F}(f) \cdot \mathcal{F}(g)^*) \tag{2}$$

where $\mathcal{F}$ and $\mathcal{F}^{-1}$ denote the forward and inverse Fourier transform, $^*$ the complex conjugate and $\cdot$ the element-wise product. pyTME is shipped with the following scoring methods.

Cross-correlation (CC):

$$\mathcal{F}^{-1}(\mathcal{F}(f) \cdot \mathcal{F}(g)^*) \tag{3}$$

Normalized cross-correlation (CORR[1,2]):

$$\frac{CC(f,g) - \overline{g} \cdot CC(f,m)}{(CC(f^2,m) - \frac{CC(f,m)^2}{N_g}) \cdot \sigma_g} \tag{4}$$

where m is masking g, $N_g$ is the number of elements in g , $\overline{g}$ is the average of g and $\sigma_g$ is the standard deviation of g.

Given the definitions of CC and CORR, they can be easily extended to other used scoring methods, such as Laplacian cross-correlation (LCC):

$$CC(\nabla^2 f, \nabla^2 g) \tag{5}$$

where $\nabla$ denotes the differential operator.

The correlation about the mean (CAM[1–3]) is defined as:

$$CORR(f, g - \overline{g}) \tag{6}$$

The fast local correlation function (FLCF[4]) used in PyTOM[5] and STOPGAP[6] is implemented as:

$$\frac{CC(f, \frac{g*m - \overline{g*m}}{\sigma_{g*m}})}{N_m * \sqrt{\frac{CC(f^2,m)}{N_m} - (\frac{CC(f,m)}{N_m})^2}} \tag{7}$$

where $N_m$ is the number of voxels within the template mask $m$.

In certain scenarios, it might be desirable to not only mask the template but also the target, e.g., when investigating membrane proteins to mask out the membrane. PyTOM[5] and STOPGAP[6] support masking the target, but the target mask is applied after score computation, which is not equivalent to applying it during score computation.[7] pyTME implements the masked cross-correlation (MCC[7]) as follows:

$$\frac{CC(f,g) - \frac{CC(f,m) \cdot CC(t,g)}{CC(t,m)}}{\sqrt{(CC(f^2,m) - \frac{CC(f,m)^2}{CC(t,m)}) \cdot (CC(t,g^2) - \frac{CC(t,g)^2}{CC(t,m)})}} \tag{8}$$

where $t$ is the mask for target $f$. In this notation, both $f$ and $g$ are already masked by $t$ and $m$, respectively.

In practice, all methods outlined above normalize the cross-correlation by factoring in the local variance of the target.

Non-exhaustive template matching can utilize the scoring methods outlined in the previous section as well as additional ones that cannot be easily evaluated through Fourier transforms. These are outlined in Vasishtan and Topf[8] and are repeated here for thoroughness.

Let $p_1, p_2, ..., p_n$ be points in the target $f$ and $q_1, q_2, ..., q_n$ be points in the template $g$, then the partial least-square (PLSQ) difference between $f$ and $g$ follows as:

$$\sum_{i=1}^{n} ||f(\mathbf{p}_i) - g(\mathbf{q}_i)||^2 \tag{9}$$

PLSQ simplifies to least-square (LSQ) if $p = q = V$, where $V$ comprises all points defining $f$.

Let $\vec{v_1}, \vec{v_2}, ..., \vec{v_3}$ and $\vec{w_1}, \vec{w_2}, ..., \vec{w_n}$ be normal vectors on the target $f$ and template $g$, then the normal vector score (NVS) follows as:

$$\frac{1}{N}\sum_{i=1}^{N} \frac{\vec{v}_i \cdot \vec{w}_i}{||\vec{v}_i||\,||\vec{w}_i||} \tag{10}$$

The Envelope Score (ENV) quantifies atom fitting within a density map and relies on the definition of a volume boundary $\rho_f$ that is used to redefine the target f so that:

$$f'(\mathbf{p}) = \begin{cases} -x, & \text{if } f(\mathbf{p}) > \rho_f \\ x, & \text{otherwise} \end{cases}$$

Based on this, the template g can be redefined so that:

$$g'(\mathbf{p}) = \begin{cases} 2, & \text{if } f'(\mathbf{p}) = -1 \\ -2, & \text{if } f'(\mathbf{p}) = 0 \\ g(\mathbf{p}), & \text{otherwise} \end{cases}$$

which yields the envelope score as:

$$\sum_{\mathbf{p} \in P} f'(\mathbf{p}) \cdot g'(\mathbf{p}) \tag{11}$$

Given points $\mathbf{x}_i$ in the target f and the set Y taken from the template g, the Chamfer distance between f and g is described as:

$$\frac{1}{|X|}\sum_{\mathbf{x}_i \in X} \inf_{\mathbf{y} \in Y} ||\mathbf{x}_i - \mathbf{y}||_2 \tag{12}$$

When considering the voxel densities of the probe f and the target g, the Mutual Information (MI) between them, given the marginal distributions p(f) and p(g) and their joint distribution p(f,g), is expressed as:

$$\sum_{f,g} p(f,g) \log \frac{p(f,g)}{p(f)p(g)} \tag{13}$$

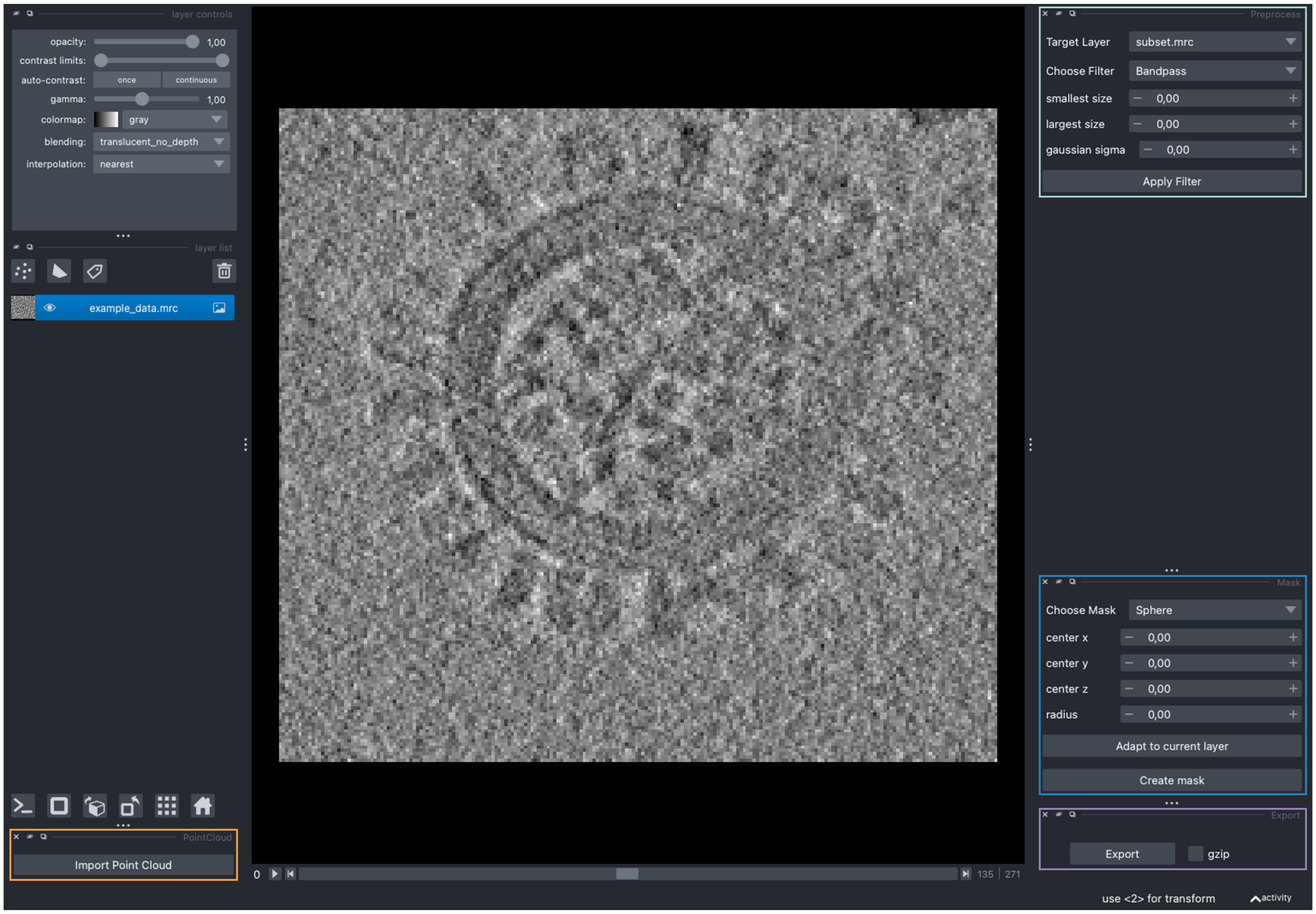


**Figure S1.** Screenshot of the Napari viewer plugin for data preprocessing. The dock menus encircled on the right side "Preprocess", "Mask" and "Export" enable users to choose from different preprocessing options and to export the final results. Similarly, the "Point-Cloud" menu encircled on the bottom left side simplifies importing, evaluating and filtering high-scoring candidate orientations.

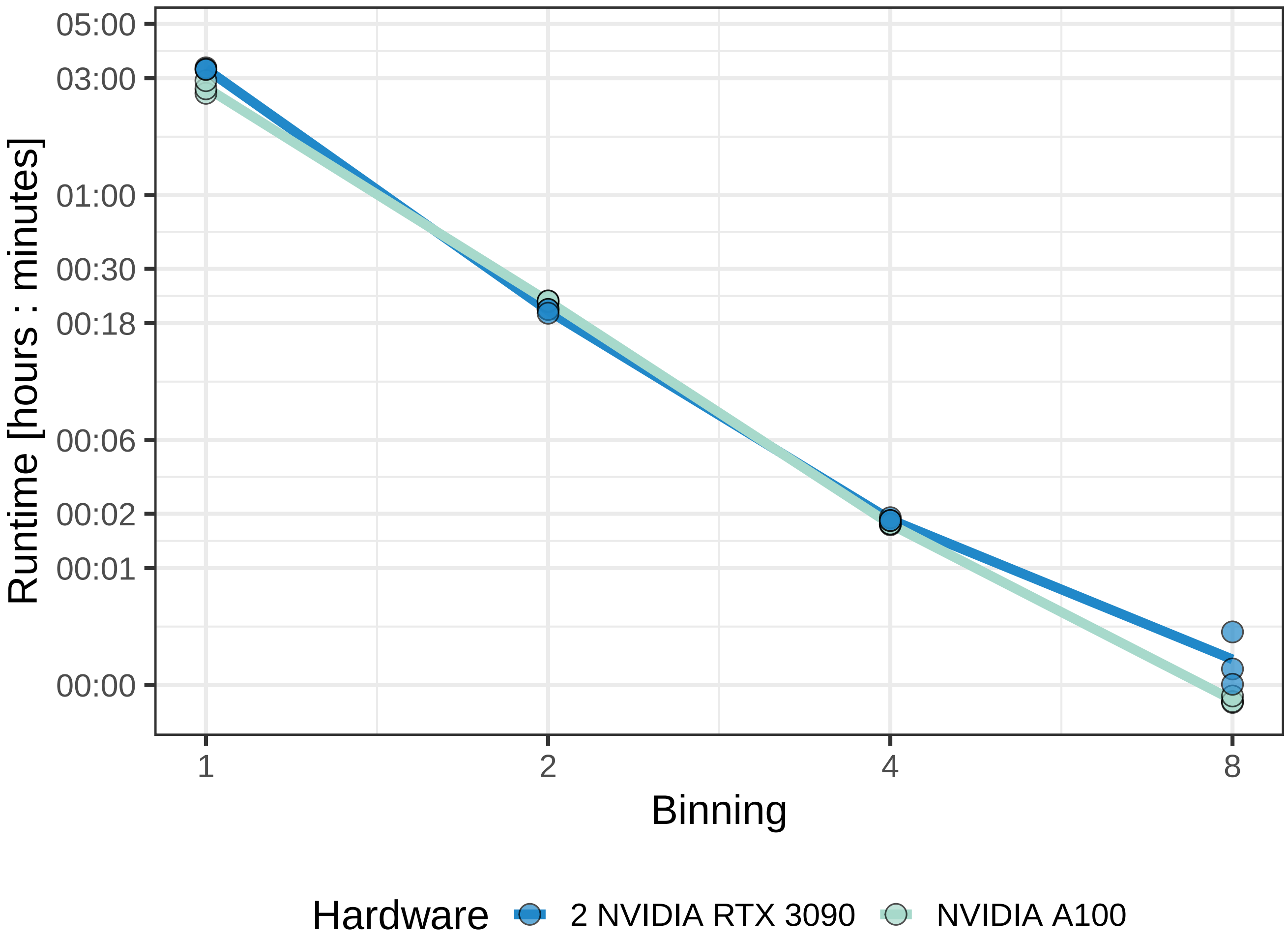


**Figure S2.** Relation between pyTME runtime and tomogram binning. At bin one, the dimension of the target was 1800, 3708, 3708, and of the template 190, 190, 190. The dimensions at bins two, four, and eight can be obtained by division of the dimensions at bin one and rounding to the next integer. Different binnings of the tomogram were obtained through reconstruction, while the template was resampled to different binnings from bin two using cubic spline interpolation. pyTME was run on an NVIDIA A100 and 2 NVIDIA RTX 3090 both sampling 1,992 rotations, computing CORR (Equation 4), without edge and Fourier padding. Each binning was run in triplicates. For bin1 pyTME was run using mixed precision and memory-mapping.
